# Preserved cortical hierarchy characterizes vision without awareness after visual cortex damage

**DOI:** 10.64898/2026.08.26.747212

**Authors:** Davide Orsenigo, Andrea I. Luppi, Matteo Diano, Tommaso Ciorli, Alessio Borriero, Matteo Panormita, Hanna E. Willis, Giovanni Petri, Holly Bridge, Marco Tamietto

## Abstract

After primary visual cortex (V1) damage, some patients successfully detect stimuli yet deny seeing them, a rare condition known as “blindsight”. Blindsight provides a unique natural experiment in consciousness research: visual information guides behavior without entering awareness. Why only some patients with comparable lesions show blindsight is unknown, and current accounts have focused on spared, V1-bypassing pathways. One extant question concerns how the functional organization of the remaining brain helps explain the divergent behavioral outcomes of V1 damage. Using resting-state fMRI, we quantified regional coupling as mutual information in the largest available cohort of psychophysically defined blindsight-positive and negative patients with age-matched controls. Despite overlapping lesions and field defects, blindsight-positive patients retained a control-like sensory–association coupling pattern, whereas blindsight-negative patients showed reduced unimodal–transmodal differentiation, reflecting lower coupling of unimodal cortex. This differentiation tracked blind-field stimulus detection and predicted performance across the patient cohort. The ipsilesional posterior thalamus, in a location consistent with the pulvinar, showed the same group difference and stronger coupling with contralesional visual and frontal pre-motor regions in blindsight-positive patients. Together, our findings outline the complementary contribution of the local and global functional organization through which V1-bypassing visual signals can influence behavior. An intrinsic cortico-subcortical functional architecture previously implicated in conscious states also bears on visual behavior without awareness.

## INTRODUCTION

Visual awareness and visually guided behavior normally feel inseparable. We see a stimulus, report seeing it, and act on it. Neurological injury can break this apparent unity. Damage to the primary visual cortex (V1) produces clinical blindness in the contralesional visual field, yet some patients still discriminate, localize, or respond to stimuli they deny seeing [1–3]. This condition, known as “blindsight”, is central to consciousness science, because it reveals that visual information can remain behaviorally effective even when it fails to enter subjective experience [4–14]. It also poses a deceptively simple question: why do some people with comparable V1 lesions retain residual visual behavior, whereas others fail to demonstrate any non-conscious response?

Classical accounts have focused on spared visual path-ways that bypass V1, documented in both human and non-human primates. Candidate routes include pathways through the superior colliculus and pulvinar, geniculo-extrastriate projections from the lateral geniculate nucleus (LGN) to the middle temporal (MT) area and other extrastriate cortices, and transcallosal connections between the intact and damaged hemispheres [15– 24]. However, identifying spared pathways does not fully explain how the signals they carry engage the wider networks supporting behavior in some patients but not in others. A signal that bypasses V1 must still engage distributed functional systems that select sensory evidence and translate it into behavior. An unresolved question is therefore whether the large-scale functional organization of the remaining brain helps explain the divergent behavioral outcomes of V1 damage.

Traditional neuroimaging approaches to visual awareness have focused on task-evoked functional magnetic resonance imaging (fMRI) to contrast events that do or do not reach subjective experience, revealing regional responses transiently associated with conscious and non-conscious visual processing [19, 25–30] Resting-state fMRI offers a complementary approach because it characterizes intrinsic network organization without requiring performance of tasks that neurological injury may compromise [31–33]. Notably, intrinsic brain dynamics can also anticipate fluctuations in visual awareness under constant stimulation [34], while resting-state measurements can predict individual patterns of task-evoked activity [35, 36], help explain the behavioral consequences of focal lesions [37–39], and predict the effects elicited by intracranial stimulation [40]. Gradient-based analyses of resting-state functional connectivity situate discrete large-scale networks along a continuum from unimodal sensory and motor cortices to transmodal association areas [41–43]. This sensory-to-association (S–A) axis is also expressed across cytoarchitecture, intracortical myelination, gene expression, receptor distributions, and maturation [43, 44], thus providing a compact coordinate system [45] for understanding how sensory signals are progressively integrated into action and cognition [46].

This intrinsic functional architecture is increasingly linked to changes in global conscious states. Anesthesia and disorders of consciousness (e.g., coma, vegetative states) have been associated with reduced differentiation between the unimodal and transmodal poles of cortical organization, alongside altered thalamocortical coordination [47–49]. In non-human primates, selective stimulation of high-order thalamic nuclei can restore behavioral arousal and wake-like cortical dynamics during ongoing anesthesia [50–52], while also reversing anesthesiainduced disruptions of distributed cortical hierarchical organization [48]. Yet these state-level findings leave open whether the same architecture also supports the behavioral use of visual information outside awareness in an otherwise conscious brain.

Blindsight provides a unique natural test of this question. Patients are awake, responsive, and globally conscious, yet visual information from a portion of their visual field is no longer experienced. Crucially, blindsight-positive (B+) and blindsight-negative (B−) patients give the same subjective report (no visual experience in the blind field), but differ in objective evidence that unseen stimuli can still guide behavior. This comparison thus shifts the question from whether a person is conscious to whether visual information remains behaviorally available without acknowledged awareness. If S–A differentiation reflects only the global conscious state, we would not expect systematic differences between B+ and B− patients, who share preserved wakefulness and report no blind-field visual experience. Conversely, if this largescale functional architecture also indexes non-conscious visual processing, B+ patients should exhibit a more preserved cortical hierarchy than B− patients, approaching the pattern observed in controls.

Here, we addressed these questions in the largest resting-state fMRI cohort of patients behaviorally characterized for blindsight. Patients were classified as B+ if they detected drifting gratings in the blind field above chance in a two-interval forced-choice task (*n* = 8), or B− if their performance did not differ significantly from chance (*n* = 8). We compared both groups with age-matched healthy controls (*n* = 17) and characterized whole-brain functional architecture using mutual information (MI), a symmetric measure of statistical dependence between regional BOLD signals [53]. We then summarized each of 360 cortical and 32 subcortical parcels by its mean MI with all other analyzed regions [54, 55] (Fig. 1). This measure therefore quantifies each region’s average coupling with the rest of the brain. We asked, across three nested levels of analysis: (i) whether each group’s intrinsic map aligns with the canonical S–A axis; (ii) whether unimodal–transmodal differentiation tracks individual blind-field detection performance; and (iii) which cortical and subcortical coupling patterns distinguish the patient groups.

**FIG. 1.**
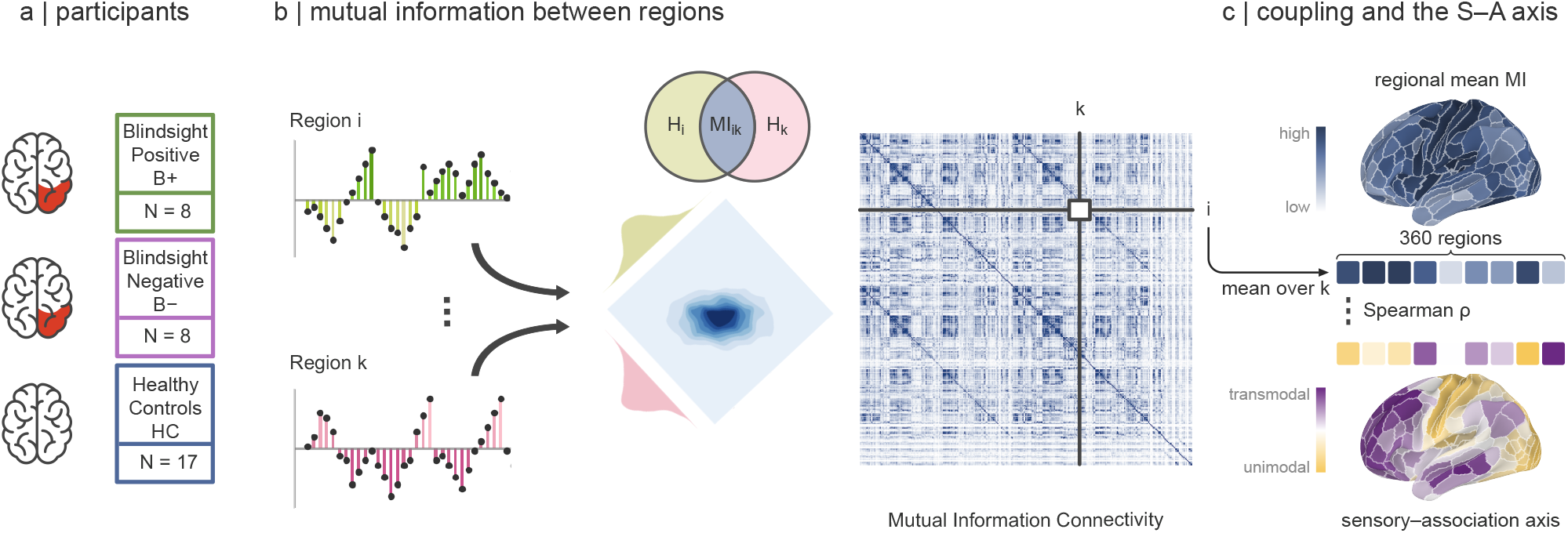
Study cohort and analysis pipeline. **a**, Participant groups. Blindsight-positive (B+, *n* = 8) and blindsight-negative (B−, *n* = 8) patients had unilateral, V1-centered lesions (red). B+ patients detected blind-field stimuli above chance, whereas B− patients performed at chance. Healthy controls (HC, *n* = 17) served as the reference group. **b**, Resting-state BOLD time series were extracted from 360 cortical and 32 subcortical parcels and *z*-scored. Pairwise mutual information (MI) was estimated using a Gaussian model, yielding a 392 *×* 392 MI matrix. The schematic illustrates the marginal and joint distributions of two regional signals *i* and *k*, and the corresponding information diagram, in which the overlap of the entropies *H*_*i*_ and *H*_*k*_ is their mutual information, MI_*ik*_. Group-level analyses used time series concatenated across participants within each group; individual-level analyses used each participant’s time series separately. **c**, For each parcel, MI was averaged over all other cortical and subcortical parcels to obtain regional mean MI (top; the strips represent the parcel-wise vectors), indexing average statistical coupling with the rest of the brain. Group-level cortical maps were correlated with the S–A axis (bottom) across the 360 cortical parcels (Spearman’s *ρ*), with significance assessed using spin permutations that preserve spatial autocorrelation. Individual maps were additionally used to quantify unimodal–transmodal differentiation. See Methods for details.

We found that B+ patients retained an S–A pattern of regional coupling resembling that of healthy controls. Conversely, B− patients showed reduced differentiation between the unimodal and transmodal poles of this axis. This reduction primarily reflected lower mean coupling of unimodal regions with the rest of the brain. Across patients, greater unimodal–transmodal differentiation was correlated with better blind-field detection performance and predicted held-out scores in within-cohort cross-validation. We further identified an ipsilesional ventral posterior thalamic territory, in a location consistent with the pulvinar, whose coupling profile distinguished B+ from B− patients. This thalamic parcel showed stronger coupling in B+ than in B− patients with three contralesional cortical regions: V1, the frontal eye field (FEF), and the dorsal visual transitional area (DVT) [54], a posteromedial region adjacent to area prostriata [56, 57].

Overall, our findings extend accounts of blindsight beyond spared visual pathways, linking residual visual behavior after V1 damage to preserved sensory-to-association organization and posterior thalamocortical coupling. More broadly, an architecture implicated in global states of consciousness is also associated with the capacity to use visual information without awareness.

## RESULTS

### Residual visual behavior diverges despite overlapping V1-centered lesions and visual-field defects

Residual visual detection was assessed at locations within each patient’s perimetrically defined blind field using a two-interval forced-choice task [18, 58, 59]. Participants maintained central fixation and indicated which of two tone-cued intervals contained a drifting achromatic Gabor patch, a design that minimizes reliance on a subjective detection criterion. Because residual vision after V1 damage is elicited most reliably by high-contrast moving stimuli [17, 18, 58, 59], performance was quantified as percentage correct on high-contrast trials (50% and 100% contrast), averaged across tested blind-field locations.

Eight patients detected stimuli significantly above the 50% chance level at all tested blind-field locations and were classified as blindsight-positive (B+; cumulative binomial test, *P <* 0.05). Eight patients whose performance did not significantly differ from chance at any tested location were classified as blindsight-negative (B−). Three additional patients who showed abovechance detection at only one of two tested locations were excluded from all analyses (see Methods). Mean detection accuracy was 88.8 *±* 7.8% in B+ and 52.5 *±* 11.3% in B− patients (mean s.d.; Fig. 2a). In the sighted hemifield, both groups performed near ceiling (B+: 91.3 *±* 8.3%; B−: 93.8 *±* 8.8%), excluding general task or attentional difficulties as the source of chance performance in B− patients. The behavioral distinction therefore concerned objective evidence of residual detection, rather than differences in reported awareness.

**FIG. 2.**
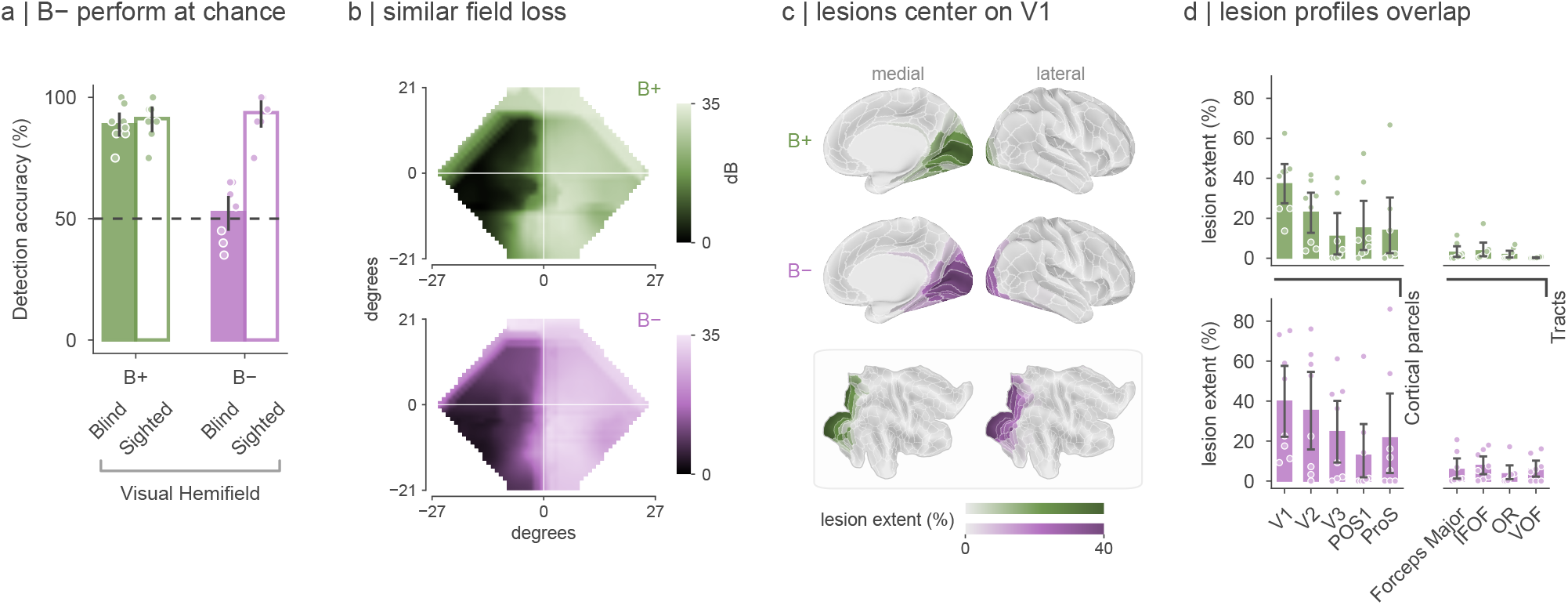
Residual visual detection diverges despite overlapping V1-centered lesions and visual-field defects. **a**, Detection accuracy of B+ (green) and B− (purple) patients in a two-interval forced-choice task with drifting Gabor patches, for high-contrast trials (50% and 100% contrast) in the blind field (filled bars) and the sighted field (open bars). Blind-field scores are averaged across tested locations, and central fixation was monitored with eye tracking. Bars show group means with 95% bootstrap confidence intervals (10,000 resamples); points show individual patients. The dashed line marks chance level (50%). B+ patients performed above chance at every tested blind-field location (cumulative binomial test, *P <* 0.05; 20 high-contrast trials per location), whereas B− patients did not exceed chance at any location. **b**, Group-averaged Humphrey visual-field perimetry maps for B+ (top, *n* = 6) and B− (bottom, *n* = 8) patients; darker shading indicates lower sensitivity (dB). Fields of patients with left-hemisphere lesions were mirrored so that the blind field appears on the left. Axes show visual-field coordinates in degrees of visual angle. **c**, Mean lesion extent of each cortical parcel (percentage of the parcel lesioned, averaged over the patients of each group) in the ipsilesional hemisphere, after left-hemisphere lesions were mirrored to the right: medial and lateral inflated views for B+ (top) and B− (middle) patients, and flat maps of the same hemisphere (bottom; B+ left, B− right). Damage is greatest in the medial occipital cortex around the calcarine sulcus. **d**, Lesion extent (percentage of each region affected) for B+ (top) and B− (bottom) patients, in the cortical parcels and white-matter tracts lesioned reliably across the 16 patients (one-sided Wilcoxon test against zero, FDR-corrected) (forceps major; IFOF, inferior fronto-occipital fasciculus; OR, optic radiation; VOF, vertical occipital fasciculus). Bars show group means with 95% bootstrap confidence intervals; points show individual patients. Lesion burden was greatest in early visual cortex and variable across extrastriate parcels, but lesion extent in no parcel or tract correlated with blind-field detection (Spearman, all *P >* 0.05, FDR-corrected; Supplementary Fig. 4).

We then compared the perimetric profiles of these behaviorally defined groups. Humphrey perimetry confirmed severe contralesional visual-field loss in both groups, with group-averaged sensitivity maps showing broadly overlapping regions of reduced sensitivity (Fig. 2b; individual fields in Supplementary Fig. 1). Because stimulus locations were selected within each patient’s perimetrically impaired field, above-chance detection in B+ patients reflected residual performance within clinically defined field defects.

To characterize the anatomical context of this dissociation, we mapped lesion topography after flipping left-hemisphere lesions so that all lesions lay in the right hemisphere by convention. In both groups, damage was centered on the medial occipital cortex around the calcarine sulcus, with greatest involvement of V1 and variable extension into adjacent early and higher-order visual areas (Fig. 2c; individual lesions in Supplementary Fig. 2). Quantification across atlas-defined cortical parcels and white-matter tracts revealed broadly overlapping lesion profiles (Fig. 2d). Total lesion volume did not differ significantly between groups (B+: 17.1 *±* 11.6 cm^3^; B−: 25.1 *±* 22.6 cm^3^; two-sided Brunner– Munzel *W* = *−*0.390, *P* = 0.704; Hedges’ *g* = *−*0.42 [CI: *−*1.36, 0.52]; Supplementary Fig. 3a). Lesion volume was not associated with detection accuracy (Spearman’s *ρ* = *−*0.31 [CI: *−*0.70, 0.23], *P* = 0.24; Supplementary Fig. 3b), and all B+ vs. B− contrasts reported below remained significant when lesion volume, or the extent of damage outside V1, was used as a covariate (Supplementary Table 1).

We further examined whether regional lesion burden was associated with residual detection across patients. Analyses included cortical parcels and white-matter tracts meeting a minimum-involvement criterion (lesion extent ≥ 10% in at least one patient). No parcel or tract showed a significant Spearman correlation between lesion extent and blind-field detection after FDR correction (all adjusted *P >* 0.05; Supplementary Fig. 4). A mixed-effects model including age and time since stroke likewise revealed no significant association between regional lesion extent and detection performance (see Methods).

Together, these analyses establish a behavioral dissociation between patients with overlapping V1-centered lesions and perimetric deficits, in whom residual detection was not significantly associated with any lesion measure examined. We next asked whether this dissociation was reflected in the intrinsic functional organization of cortical and subcortical networks.

### Cortical coupling aligns with the S–A axis in blindsight-positive patients

We next examined whether the cortical distribution of intrinsic coupling in the two behavioral groups aligned with the independently defined S–A axis [43], using healthy controls as a comparison group.

Within each group (healthy controls, HC; B+; B−), we concatenated regional BOLD time series across participants and estimated pairwise MI between all analyzed cortical and subcortical parcels. MI quantifies how much knowing one region’s activity reduces uncertainty about another’s, providing a measure of statistical dependence. For each cortical parcel, we averaged its MI with all other analyzed parcels, yielding a cortical map of mean coupling for each group (Fig. 3a).

**FIG. 3.**
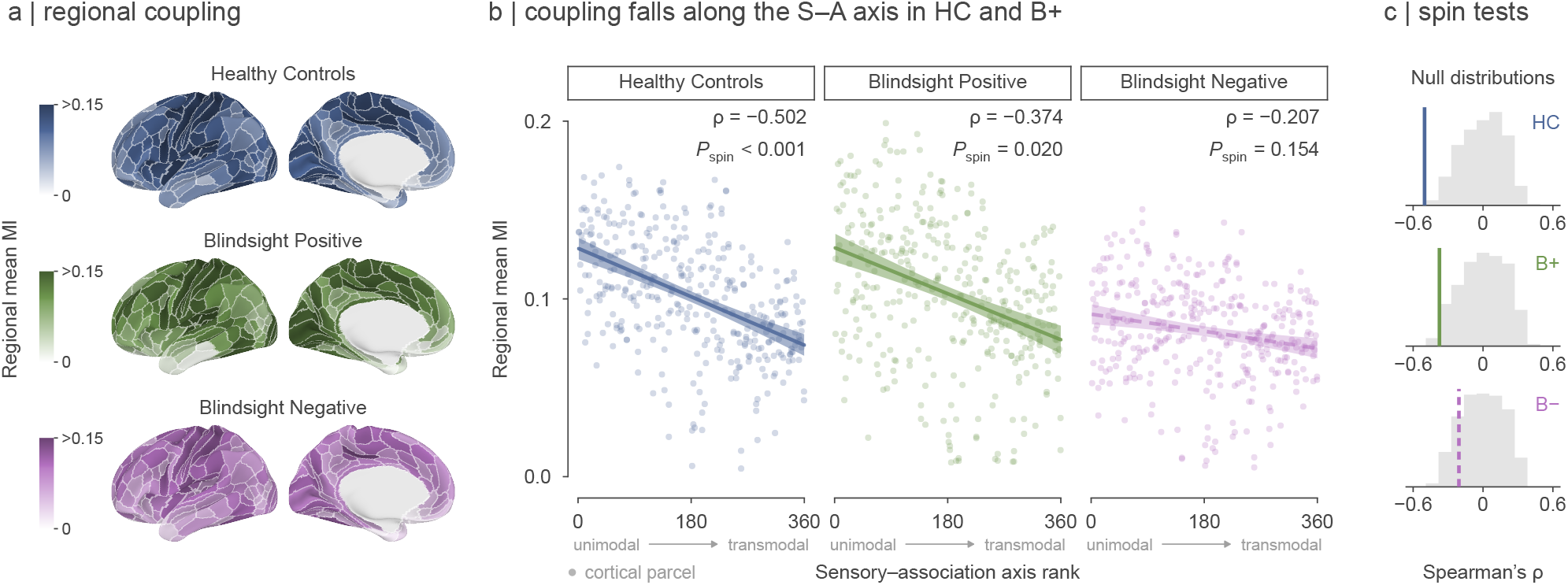
Cortical coupling aligns with the S–A axis in controls and blindsight-positive patients. **a**, Group-level cortical maps of regional mean mutual information (MI) for healthy controls (blue), blindsight-positive patients (green) and blindsight-negative patients (purple). Maps show the contralesional (left) hemisphere in lateral (left) and medial (right) views. For each group, regional BOLD time series were *z*-scored within each participant and concatenated across participants. Pairwise MI was estimated among all 392 parcels, and each cortical parcel’s MI was averaged over all other parcels. Darker shading indicates higher mean MI. Color scales are specific to each group but share the same limits; values above 0.15 saturate. **b**, Regional mean MI against each parcel’s rank on the S–A axis among the 360 cortical parcels, from unimodal sensorimotor cortex (low ranks) to transmodal association cortex (high ranks), as marked under each panel. The axis gives homologous parcels of the two hemispheres the same position, so each such pair shares a rank. Each point is one cortical parcel. Lines and shaded bands show linear fits with 95% confidence intervals, for display only; the dashed line marks a non-significant correlation. Alignment was quantified with Spearman correlations across the 360 cortical parcels (values in each panel). **c**, Null distributions (gray) of the map–axis Spearman correlation, from 5,000 spin permutations that preserve spatial autocorrelation. Colored lines mark the observed correlations (dashed: non-significant); rows follow the group order in panel a. These tests assess each map’s alignment with the S–A axis; differences between groups are tested in Fig. 4.

We assessed the alignment of these group-level maps with the S–A axis, oriented from unimodal sensorimotor to transmodal association cortex. For each group, we computed the Spearman correlation between regional mean MI and S–A position across the 360 cortical parcels. Significance was evaluated against null distributions from spin permutations that preserve cortical spatial autocorrelation (see Methods).

In healthy controls, mean coupling was highest in unimodal sensory and motor cortices and decreased toward transmodal association cortex (Fig. 3a,b), yielding a significant negative correlation with S–A position (Spearman’s *ρ* = *−*0.502, *P*_spin_ *<* 0.001; Fig. 3c).

B+ patients showed a similar organization. Their mean-coupling map retained a significant negative correlation with the S–A position (*ρ* = *−*0.374, *P*_spin_ = 0.020), despite unilateral V1 damage and the absence of reported blind-field visual experience. In B− patients, the map– axis correlation was also negative but numerically weaker and did not reach significance against the spatial null distribution (*ρ* = *−*0.207, *P*_spin_ = 0.154; Fig. 3b,c).

The magnitude of the negative map–axis correlation therefore decreased progressively from HC to B+ to B−. Because these spin tests assess each map’s alignment with the axis rather than differences between groups, we next tested whether this pattern was expressed in individual unimodal–transmodal coupling differences and whether those differences were associated with residual blind-field detection.

### Unimodal–transmodal differentiation tracks residual visual behavior

We next tested whether the group-level pattern was expressed in individual participants and related to residual visual performance. For each participant separately, we computed regional mean MI (each parcel’s average coupling with all other cortical parcels) and averaged these values across parcels assigned to unimodal (visual and somatomotor) or transmodal networks (frontoparietal and default mode) [60] (Fig. 4a). Subtracting the transmodal from the unimodal average provided an individual measure of coupling differentiation between the two poles of the S–A axis, complementing the map–axis correlations reported above (Fig. 3b).

**FIG. 4.**
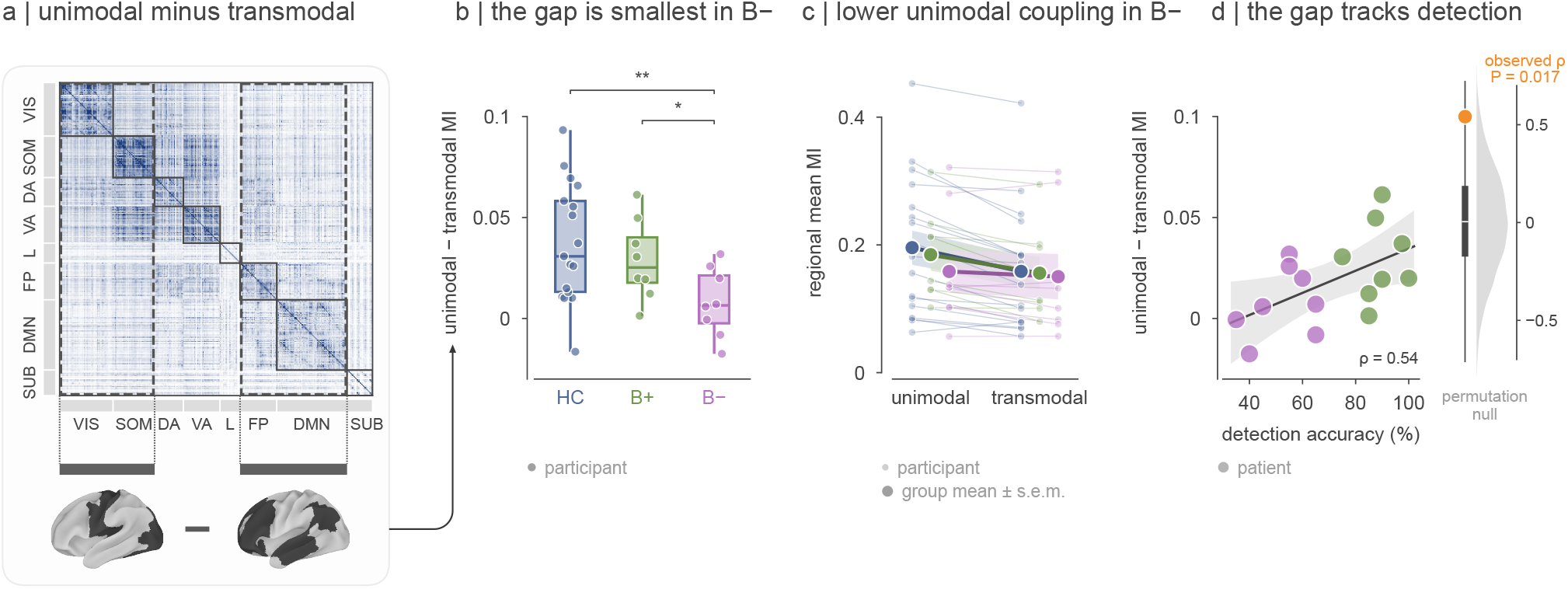
Unimodal–transmodal coupling differentiation tracks residual visual detection. **a**, Computation of participant-level coupling differentiation from the mutual information (MI) matrix, ordered by resting-state network (healthy controls shown). Regional mean MI is each cortical parcel’s average MI with all other cortical parcels. Regional means were averaged across parcels in unimodal networks (visual and somatomotor) and in transmodal networks (frontoparietal and default mode). Differentiation was defined as the unimodal minus the transmodal average. Dashed rectangles mark the matrix columns contributing to each average; cortical masks show the corresponding networks. VIS, visual; SOM, somatomotor; DA, dorsal attention; VA, ventral attention; L, limbic; FP, frontoparietal; DMN, default mode; SUB, subcortical. **b**, Unimodal–transmodal differentiation in healthy controls (HC), B+, and B−. Points show individual participants; boxes show the median and interquartile range, and whiskers the most extreme values within 1.5 times the interquartile range. Differentiation was greater in HC and B+ than in B− patients (one-tailed Brunner–Munzel tests, FDR-corrected: HC vs. B−, *P* = 0.005; B+ vs. B−, *P* = 0.025). HC and B+ did not differ significantly (*P* = 0.251). ^*^*P <* 0.05, ^**^*P <* 0.01. **c**, Regional mean MI averaged over unimodal and over transmodal parcels for each participant (small points, joined by thin lines), colored by group as in b. Large points joined by lines show the group means; shaded bands show the s.e.m. **d**, Unimodal–transmodal differentiation against blind-field detection accuracy across the 16 classified patients (Spearman’s *ρ* = 0.541 [CI: 0.12, 1], permutation *P* = 0.017). Points show individual patients, colored by group as in b. The line and gray band show a linear fit with its 95% confidence interval, for display only. Right, the null distribution of *ρ* from 10,000 permutations of detection scores across patients (gray density and box plot); the orange point marks the observed *ρ*.

Both B+ patients (Brunner–Munzel *W* = 2.358, *P* = 0.025, Hedges’ *g* = 1.069 [CI: 0.21, ∞)) and healthy controls (*W* = 3.463, *P* = 0.005, *g* = 1.073 [CI: 0.33, ∞)) showed greater unimodal–transmodal differentiation than B− patients (one-tailed tests, FDR-corrected; Fig. 4b), whereas HC and B+ patients did not differ significantly (*W* = 0.683, *P* = 0.251, *g* = 0.297 [CI: *−*0.39, ∞)). The reduced differentiation suggested by the B− group map was thus also evident at the participant level.

The smaller gap in B− patients primarily reflected lower mean coupling of unimodal regions with the rest of the brain, whereas transmodal averages were more similar across groups (difference between the B+ and B− group means: unimodal 0.026, transmodal 0.006; Fig. 4c). Neither average differed significantly between B+ and B− patients on its own (unimodal: *W* = 0.595, *P* = 0.562, *g* = 0.302 [CI: *−*0.64, 1.23]; transmodal: *W* = 0.488, *P* = 0.636, *g* = 0.059 [CI: *−*0.87, 0.98]; two-tailed Brunner–Munzel): the group difference lay in the within-participant contrast between the two poles. Notably, group differences persisted when the unimodal average was restricted to bilateral somatomotor cortex (B+ vs. B−: *W* = 2.820, *P* = 0.007, *g* = 1.039 [CI: 0.18, ∞); HC vs. B−: *W* = 3.174, *P* = 0.003, *g* = 0.991 [CI: 0.26, ∞); one-tailed Brunner–Munzel), indicating that the reduced coupling differentiation in B− patients was not confined to the visual cortex.

A mixed-effects model of regional MI tested group differences in the unimodal–transmodal contrast, with participant identity as a random effect. The model confirmed greater differentiation in HC and B+ than in B− patients (HC vs. B−: *Z* = 7.917, *P <* 0.001, *b* = 0.0291 [CI: 0.0219, 0.0363]; B+ vs. B−: *Z* = 4.869, *P <* 0.001, *b* = 0.0209 [CI: 0.0125, 0.0293]). It also detected an HC–B+ difference (*Z* = 2.239, *P* = 0.025, *b* = 0.0082 [CI: 0.0010, 0.0155]), which was not significant in the participant-level Brunner–Munzel comparison. Because this model treats each parcel’s value as a separate observation, we also fitted a participant-specific random slope for the unimodal–transmodal gap in the patient groups; the B+ vs. B− difference remained significant (*Z* = 2.26, *P* = 0.024, *b* = 0.0209 [CI: 0.0028, 0.0390]), also when lesion volume, age, sex, time since stroke and lesion laterality were allowed to modulate the gap (*Z* = 2.35, *P* = 0.019, *b* = 0.0203 [CI: 0.0034, 0.0373]; Supplementary Results).

We then examined the behavioral relevance of this individual measure, testing its association with blind-field detection accuracy. Across the 16 patients, greater unimodal–transmodal differentiation was associated with higher detection accuracy (Spearman’s *ρ* = 0.541 [CI: 0.12, 1], permutation *P* = 0.017; Fig. 4d). Because the same detection scores also defined B+ and B− status, this correlation characterizes the continuous behavioral variation underlying the group distinction, rather than providing an independent validation of it. The association remained after adjustment for age, sex, time since stroke and lesion laterality (partial *ρ* = 0.619 [CI: 0.12, 1], *P* = 0.017; Supplementary Fig. 5e).

To assess predictive performance beyond this in-sample association, we used leave-one-out cross-validation. The individual unimodal–transmodal difference predicted held-out detection scores with a cross-validated *R*^2^ of 0.15 (permutation *P* = 0.025). This result supports the predictive relevance of coupling differentiation within the present cohort.

Together, these analyses link residual visual behavior to the differentiation of intrinsic coupling between uni-modal and transmodal systems. We next asked which cortical and subcortical coupling profiles were associated with this behaviorally relevant pattern.

### Thalamocortical coupling distinguishes blindsight-positive from blindsight-negative patients

Having linked unimodal–transmodal differentiation to residual visual behavior, we next examined which regional coupling profiles distinguished B+ from B− patients. We focused first on the thalamus, given its role in the V1-bypassing pathways implicated in blind-sight [17, 18, 20, 61, 62]. For each participant and each of the eight thalamic parcels defined by Tian et al. [55] (four per hemisphere), we calculated the difference between the parcel’s mean MI with unimodal cortex and its mean MI with transmodal cortex. This measure indexes how strongly each thalamic parcel’s coupling differentiates the two cortical poles [44], rather than its overall coupling strength.

Among the eight parcels, only the ipsilesional ventral posterior thalamus, in a location consistent with the pulvinar, showed significantly greater unimodal–transmodal differentiation in both B+ patients and healthy controls relative to B− patients (B+ vs. B−: Brunner–Munzel *W* = 3.818, *P* = 0.008, Hedges’ *g* = 1.332 [CI: 0.43, ∞); HC vs. B−: *W* = 3.697, *P* = 0.005, *g* = 0.925 [CI: 0.20, ∞); one-tailed tests, FDR-corrected) (Fig. 5a). Across patients, greater differentiation of this parcel’s thalam-ocortical coupling was associated with higher blind-field Gabor detection accuracy (Spearman’s *ρ* = 0.555 [CI: 0.13, 1], permutation *P* = 0.015).

**FIG. 5.**
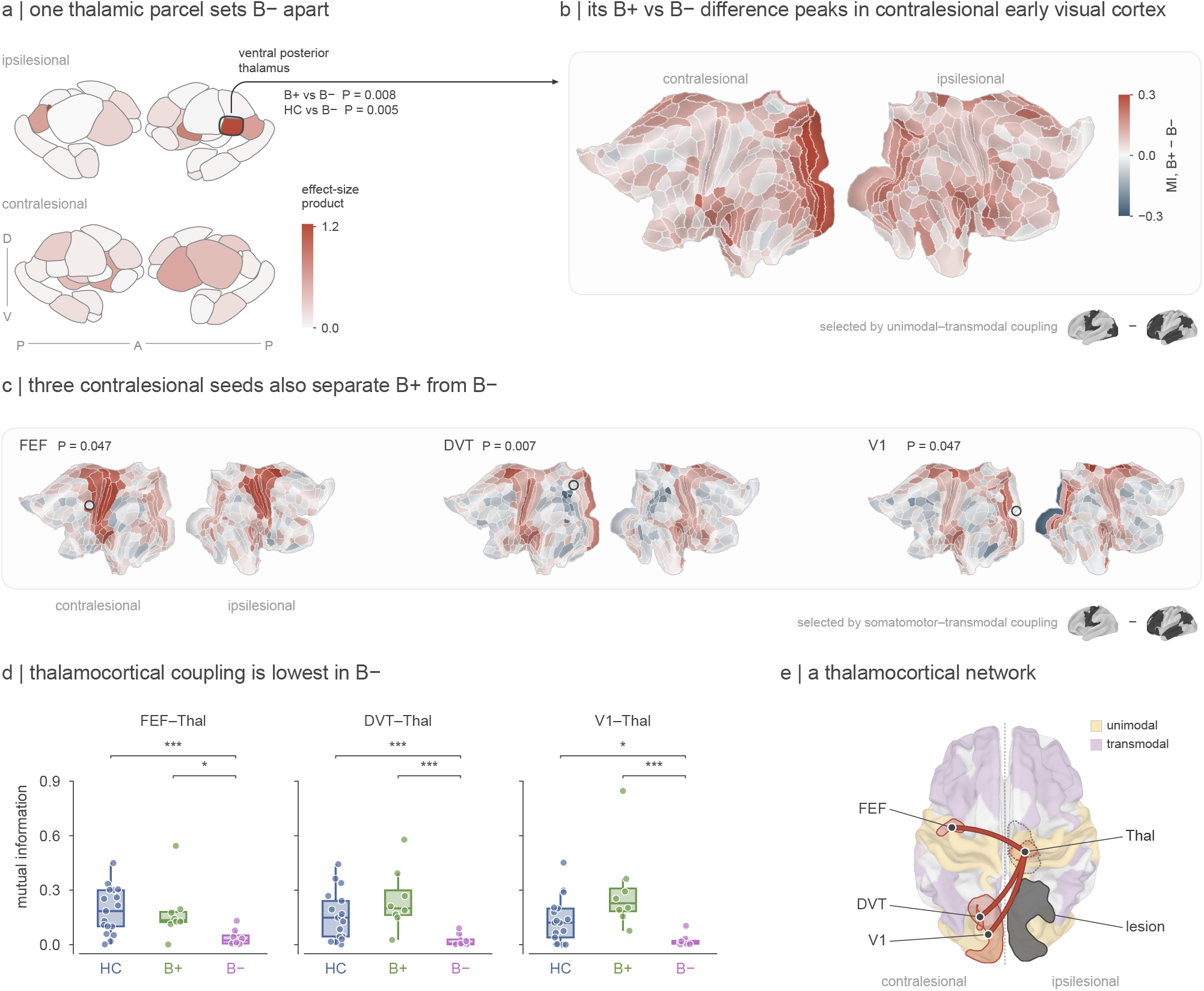
Thalamocortical coupling distinguishes blindsight-positive from blindsight-negative patients. **a**, Thalamic coupling differentiation, computed per participant as a thalamic parcel’s mean MI with unimodal cortex (visual and somatomotor networks) minus its mean MI with transmodal cortex (frontoparietal and default mode networks), shown for every subcortical parcel of the ipsilesional (top) and contralesional (bottom) hemisphere, viewed from the right (left drawing) and from the left (right drawing); A, P, D and V mark the anterior, posterior, dorsal and ventral directions; drawings adapted from the subcortex_visualization toolbox [63]. Color encodes the absolute value of the product of Hedges’ *g* for the HC vs. B− and B+ vs. B− comparisons. The outline marks the ipsilesional ventral posterior parcel (THA-VP), the only one of the eight thalamic parcels with significantly greater differentiation in both B+ patients and HC than in B− patients (one-tailed Brunner– Munzel, FDR-corrected; *P* values beside the parcel). **b**, B+ vs. B− difference in pairwise MI between the parcel identified in a and each cortical parcel, for the contralesional (left) and ipsilesional (right) hemispheres (flat maps). Red, greater coupling in B+; blue, greater in B−. The largest differences are in contralesional early visual cortex (V1–V3). Color shows the magnitude of the difference. The inset shows the networks on which the seed was selected (unimodal minus transmodal). **c**, The same maps for three contralesional seeds identified by a whole-brain analysis of somatomotor–transmodal coupling differentiation (inset). Circles mark seed locations; the color scale is as in b. The *P* values refer to the seed-level contrast (one-tailed Brunner–Munzel, FDR-corrected). **d**, MI between the ipsilesional ventral posterior thalamic parcel (Thal) and contralesional FEF, DVT and V1 for HC (blue), B+ (green) and B− (purple). Points, individual participants; boxes, median and interquartile range; whiskers, the most extreme values within 1.5 times the interquartile range. Brackets, one-tailed Brunner–Munzel comparisons against B−, FDR-corrected (^*^*P <* 0.05, ^***^*P <* 0.001). **e**, Schematic of the candidate functional network, viewed from above with the ipsilesional hemisphere on the right. Lines connect the ipsilesional ventral posterior parcel with contralesional V1, DVT and FEF, the three edges with greater coupling in B+ than in B− patients. The thalamus and its ventral posterior parcel (dashed outlines) lie beneath the cortex. Shading indicates unimodal (yellow) and transmodal (purple) cortex; the lesion is shown schematically in gray (ipsilesional parcels with mean lesion extent above 20%).

Mapping the cortical distribution of B+ vs. B− differences in coupling with the ipsilesional thalamic parcel revealed the largest differences in contralesional early visual cortex (V1–V3) (Fig. 5b). The preceding cortex-wide analysis also identified reduced somatomotor–transmodal coupling differentiation in B− patients (see above). We therefore broadened the analysis beyond the thalamus, asking which regions showed group differences in their coupling with this non-visual unimodal system relative to transmodal cortex. In this whole-brain analysis, we treated each of the 392 cortical and subcortical parcels in turn as a seed. For each participant, we calculated each seed’s mean MI with somatomotor cortex minus its mean MI with transmodal cortex, then compared these seed-specific coupling differences between B+ and B− patients.

Three cortical seeds showed greater somatomotor– transmodal differentiation in B+ than in B− patients after FDR correction across all 392 parcels, all in the contralesional hemisphere: V1 (*W* = 4.243, *P* = 0.047, *g* = 1.327 [CI: 0.43, ∞)), the frontal eye field (FEF; *W* = 4.243, *P* = 0.047, *g* = 1.520 [CI: 0.59, ∞)), and the dorsal visual transitional area (DVT; *W* = 5.580, *P* = 0.007, *g* = 1.576 [CI: 0.64, ∞)) (Fig. 5c), a parcel adjacent to area prostriata [54, 56, 57]. Contralesional V1 thus emerged from two analyses based on different measures, and contralesional frontal oculomotor (FEF) and posteromedial visual (DVT) regions also showed somatomotor–transmodal coupling profiles that distinguished the patient groups.

We then asked whether these cortical candidates were themselves differentially coupled to the ipsilesional ventral posterior thalamus. Pairwise MI between this region and each of the contralesional cortical areas was greater in B+ than in B− patients: V1 (*W* = 21.920, *P <* 0.001, Hedges’ *g* = 1.524 [CI: 0.59, ∞)), FEF (*W* = 2.541, *P* = 0.021, *g* = 1.127 [CI: 0.26, ∞)), and DVT (*W* = 12.550, *P <* 0.001, *g* = 1.737 [CI: 0.77, ∞); one-tailed Brunner– Munzel tests, FDR-corrected) (Fig. 5d). For all three edges, coupling in B− patients was also reduced relative to HC (V1: *W* = 2.417, *P* = 0.018, *g* = 1.028 [CI: 0.29, ∞); FEF: *W* = 4.368, *P <* 0.001, *g* = 1.301 [CI: 0.54, ∞); DVT: *W* = 4.229, *P <* 0.001, *g* = 1.141 [CI: 0.39, ∞)).

Comparing B+ patients with healthy controls showed that this network is not simply restored to the normal range: coupling between the ipsilesional ventral posterior thalamus and contralesional V1 was higher in B+ than in HC (two-sided Brunner–Munzel *W* = 2.814, *P* = 0.032, Hedges’ *g* = 0.930 [CI: 0.07, 1.78]; FDR-corrected), whereas the FEF and DVT edges did not differ from controls (FEF: *W* = *−*0.281, *P* = 0.78, *g* = *−*0.060 [CI: *−*0.87, 0.75]; DVT: *W* = 1.501, *P* = 0.23, *g* = 0.594 [CI: *−*0.24, 1.42]). Supra-normal thalamus–V1 coupling in the group that retains residual detection suggests compensation beyond restoration of the normal profile.

Two additional analyses characterized the anatomical specificity of this thalamic finding. First, the homologous contralesional ventral posterior parcel showed no significant B+ vs. B− difference in coupling with any of the three cortical regions (all *P >* 0.05), indicating that the effect was lateralized to the damaged hemisphere. Second, when all 392 cortical and subcortical parcels were ranked by the separation between patient groups in their coupling with these cortical regions, the ipsilesional ventral posterior thalamus ranked first under each of three criteria (rank 1 of 392): (i) the mean absolute effect size of the B+ vs. B− contrast across the circuit edges, (ii) the mean product of the HC vs. B− and B+ vs. B− effect sizes, and (iii) the Stouffer-combined score for the one-tailed B+ *>* B− test across those edges. This parcel did not overlap the lesion in any patient, and its B+ vs. B− difference held with lesion volume as a covariate (Supplementary Table 1).

Together, these analyses identified a candidate functional thalamocortical network linking the ipsilesional ventral posterior thalamus with contralesional visual (V1, DVT) and frontal (FEF) cortex (Fig. 5e). Coupling between this thalamic parcel and all three cortical regions was stronger in B+ than in B− patients, complementing the cortex-wide association between residual behavior and unimodal–transmodal differentiation.

### Robustness and limitations

We assessed whether the principal findings depended on the coupling estimator, the cortical parcellation or the hierarchy reference (Supplementary Results). The group ordering, direction of the brain–behavior association and the coupling differences involving the ipsilesional ventral posterior thalamus were reproduced with rank-Gaussian copula and *k*-nearest-neighbor MI estimators, including after a circular-shift control that preserves each signal’s temporal autocorrelation (Supplementary Table 2; Supplementary Figs. 6–9). The cortical findings were also reproduced with the 200-parcel Schaefer atlas [64] (Supplementary Fig. 10). Moreover, replacing the S–A axis with cortical T1w/T2w ratios, a structural proxy for cortical hierarchy derived independently of the functional data, again yielded significant map–hierarchy associations in HC and B+, but not in B− patients [65, 66] (Supplementary Fig. 11). Conventional functional connectivity did not reproduce the full group-discrimination pattern in the implementations tested (Supplementary Figs. 12, 13). This suggests that sensitivity to the group differences depends in part on how coupling is summarized: the Gaussian MI estimator disregards correlation sign and nonlinearly emphasizes stronger correlations before regional averaging (Supplementary Results).

Because each patient group comprised eight participants, we asked whether the principal contrasts depended on particular individuals, using participant-level resampling. The HC vs. B− and B+ vs. B− contrasts in individual unimodal–transmodal differentiation (Fig. 4) remained significant after omitting any single participant and under permutation of group labels. In an exploratory whole-pipeline relabeling analysis, the specified joint pattern of group differences and map–axis alignment recurred in only seven of 10,000 pseudo-group reassignments (empirical *P* = 8 *×* 10^*™*4^; Supplementary Fig. 14). The individual estimates used in brain–behavior analyses showed within-run consistency (intraclass correlation = 0.76; Supplementary Fig. 15). The cortical pattern was also broadly reproduced in separate analyses of each hemisphere (Supplementary Table 3; Supplementary Figs. 16, 17), and the principal contrasts persisted after adjustment for demographic and clinical covariates (Supplementary Table 1; Supplementary Fig. 5e).

Because lesioned tissue carries little shared signal and the lesions lay mostly within the visual network, part of the unimodal pole, we also asked whether the cortical findings reflected the lesions themselves. After removal of the 32 cortical parcels lesioned by at least 10% in any patient, both as targets and as sources of each parcel’s regional mean MI, the map–axis alignment remained significant in HC and B+ but not in B−, and both participant-level contrasts and the association with detection accuracy were reproduced (Supplementary Table 4; Supplementary Fig. 3).

Together, these checks indicate that the preserved versus flattened hierarchy distinguishing B+ from B− patients persists across the analytical specifications tested and is a property of the data rather than of the estimator, parcellation, hierarchy reference or any individual participant.

## DISCUSSION

Residual visual detection after V1 damage was associated with the intrinsic organization of distributed cortical and thalamocortical systems. Despite overlapping lesions and visual-field defects, B+ patients retained a sensory-to-association (S–A) coupling pattern resembling that of healthy controls, whereas B− patients showed reduced unimodal–transmodal differentiation. At the participant level, coupling differentiation tracked blind-field detection and predicted held-out performance within the cohort. At the regional level, B+ patients showed stronger coupling between an ipsilesional posterior thalamic territory and contralesional visual and frontal regions. These findings extend pathway-based accounts of blindsight by linking the behavioral use of residual visual information to the intrinsic organization of surviving brain systems. The findings extend the relevance of S–A organization beyond global changes in conscious state. Alterations in cortical functional gradients accompany anesthesia and disorders of consciousness, and central thalamic stimulation can reverse distributed hierarchical disruption while restoring behavioral arousal in anesthetized macaques [47, 48]. Complementary TMS–EEG evidence shows that cortical responses propagate across distributed regions during wakefulness but become local and short-lived during non-REM sleep [67]. Here, both patient groups were awake and responsive and reported no blind-field visual experience, yet differed in objective detection. The comparison therefore concerns the behavioral availability of visual information within a preserved conscious state, not a transition between consciousness and unconsciousness. Although our coupling measures and previous gradient analyses capture different aspects of cortical organization, their convergence suggests that an architecture implicated in conscious states also supports processing that remains effective without visual awareness.

Control-like intrinsic organization should not be equated with conscious processing of blind-field stimuli, nor are the structures involved dedicated exclusively to non-conscious processing. Indeed, preserved intrinsic organization need not imply preservation of the stimulus-evoked recurrent interactions proposed to support conscious vision [68]. In healthy observers, recognized and unrecognized objects recruit overlapping distributed networks, with changes in the strength and decodability of the responses within them [69, 70]. In the thalamus, pulvinar firing varies with stimulus visibility, and thalamo-cortical interactions track conscious perception [71–73], while pulvinar inactivation alters confidence without significantly impairing categorization accuracy [74]. What matters is likely the specific interaction, not the structure [75]. These prior studies asked which interactions accompany successful perception, whereas our comparison identifies the interactions that track whether an unseen stimulus still guides behavior despite a shared absence of reported blind-field experience.

Resting-state measurements characterize this organization without requiring performance of a visual task. Convergent evidence links intrinsic activity to behavior. For example, precuneus–visual interactions anticipate perceptual transitions during binocular rivalry [34]. Task-free measurements predict individual patterns of task-evoked activity [35], and spontaneous motor activity resembles patterns evoked by familiar movements [76]. Such findings support accounts in which spontaneous activity reflects processing patterns shaped by experience and behavioral demands [36, 77]. Our results extend this perspective to focal visual injury, suggesting that intrinsic coupling indexes the availability of coordinated processing for the use of residual sensory evidence.

The posterior thalamus anchors this distributed cortical pattern. The ipsilesional ventral posterior thalamic parcel, in a location consistent with the pulvinar, showed greater unimodal–transmodal coupling differentiation in B+ than in B− patients, and stronger coupling with the contralesional cortical regions that distinguished the groups. This finding complements lesion-network evidence implicating connectivity with the ipsilesional medial pulvinar in blindsight [23], although the parcellation does not resolve individual pulvinar nuclei. Causal evidence indicates how pulvinar circuitry could contribute. Early pulvinar lesions disrupt dorsal-stream development and later visually guided behavior in marmosets [78]. In macaques with intact V1, pulvinar inactivation reduces visual responsiveness, attentional modulation and interareal coordination, while increasing low-frequency cortical activity [79]. In V1-lesioned macaques, inactivation of the pulvinar or of its collicular input impairs visually guided saccades into the affected field [20]. These findings motivate the hypothesis that posterior thalamocortical interactions support residual vision not only by transmitting sensory signals but also by maintaining the cortical responsiveness and coordination needed to use them.

The cortical regions coupled to this thalamic territory point to selection and action as candidate contributors to residual detection. Spatial attention can improve processing of blind-field stimuli without restoring visual awareness, demonstrating that attentional selection and conscious access can dissociate after V1 damage [80]. Contralesional V1, FEF and DVT distinguished the groups through their somatomotor–transmodal coupling profiles, showing that the association with residual vision extends beyond visual systems. The behavioral contribution of a sensory region cannot be inferred solely from the information it encodes. In mice, unilateral S1 inactivation shifts tactile decision bias through superiorcolliculus-dependent circuitry rather than consistently reducing sensitivity [81]. Our two-interval forced-choice task reduces dependence on a yes/no criterion, but success still requires selecting and using sensory evidence. The somatomotor associations therefore motivate testing whether behavioral readout contributes to group differences alongside residual sensitivity. FEF offers a plausible interface between visual processing and orienting, consistent with evidence that its perturbation alters detection [82, 83]. DVT is adjacent to area prostriata, whose fast-motion responses and retinotopically organized thalamic connections warrant investigation of posteromedial visual cortex in residual vision [56, 84]. The contralesional location of all three nodes is consistent with interhemispheric contributions to post-stroke behavior and to residual visuomotor function [19, 37]. Posterior thalamus–V1 coupling was greater in B+ patients than in controls. Thus, control-like cortical organization coexisted with enhanced regional coupling, a pattern compatible with compensatory reweighting.

The present findings identify testable directions for rehabilitation research. Visual cortical activity and resting precuneus connectivity have been linked to training outcome after occipital injury [85–87]. Prospective studies could test whether unimodal–transmodal differentiation predicts training benefit beyond baseline performance and lesion characteristics, and whether it changes with improvement. Manipulating spatial certainty, distractor competition and response demands would help distinguish residual sensitivity from the selection and use of visual evidence. FEF perturbation offers a causal test of the proposed contribution to behavioral readout, given its effects on detection and awareness in neurologically intact observers [82, 83, 88, 89]. A recent report that stimulation targeting the human pulvinar in a patient with drug-resistant epilepsy elicited spatially distinct visual percepts and modulated distributed cortical activity provides a complementary rationale for investigating thalamocortical mechanisms [90]. Measuring detection, confidence and awareness separately would determine whether these interventions improve the use of unseen information, restore visual experience, or both [91].

Together, these results link residual vision to the functional organization of surviving systems at both regional and cortex-wide scales. Spared pathways constrain where visual signals can still travel. S–A coupling differentiation and posterior thalamocortical interactions identify intrinsic network features associated with their behavioral use. An architecture previously implicated in global conscious state thus also bears on whether visual information can guide behavior without reported awareness.

## MATERIALS AND METHODS

### Participants

Nineteen patients with chronic unilateral stroke involving V1 were recruited (5 women; age 24–74 years, mean *±* s.d. 50.2 *±* 15.6; time since stroke ≥ 6 months, mean 43.4*±*63.8 months; recruitment as in Willis et al. [58, 59]). Inclusion required a homonymous visual-field defect on Humphrey perimetry, a lesion confined to one hemisphere with negligible involvement of higher-order visual cortex, and no cognitive, psychiatric or motor impairment, ocular disease or other post-stroke deficit. Seventeen healthy controls were matched for age and sex (6 women; 50.2*±*23.3 years). All participants gave written informed consent. The study was approved by the University of Oxford Central University Research Ethics Committee (R60132/RE001) and conducted in accordance with the Declaration of Helsinki. Participants abstained from caffeine and alcohol for 48 h before scanning.

### Behavioral assessment and classification

Residual vision was assessed with a two-interval forced-choice contrast-detection task (for details, see Willis et al. [58]). On each trial, two tone-cued intervals were presented and a drifting achromatic Gabor patch (diameter 5^*°*^, *σ* = 0.8^*°*^, 1 cycle per degree, 10 Hz, 500 ms; background 16.5 cd m^*™*2^; viewing distance 42 cm) appeared in one of them at 1, 5, 10, 50 or 100% contrast. Participants reported which interval contained the stimulus. Stimuli were presented at two locations inside the perimetrically defined blind field and at mirror-symmetric locations in the sighted field, 20 high-contrast trials per location. Participants fixated a central cross, and fixation was monitored with an EyeLink 1000 (SR Research). Absence of visual experience in the blind field was established by verbal report after each block. Because residual vision is elicited most reliably by high-contrast moving stimuli [17, 58], the behavioral score was percent correct on 50% and 100% contrast trials, averaged across tested blind-field locations (cohort mean 70.7*±*19.1%, range 35– 100%). Patients performing above chance at every tested blind-field location (cumulative binomial test, *P <* 0.05) were classified as blindsight-positive (B+, *n* = 8); patients at chance at every location as blindsight-negative (B−, *n* = 8). Three patients who exceeded chance at one location but not the other were excluded from all analyses.

### Lesion mapping and lesion-control analyses

Lesions were manually delineated on the T1-weighted image and independently verified. Left-hemisphere lesions and the corresponding imaging data were mirrored so that all lesions lay in the right hemisphere. In controls, the right hemisphere was treated as ipsilesional and the left as contralesional throughout. Lesion extent was quantified as the percentage of voxels damaged in each Glasser parcel and in four white-matter tracts (forceps major, inferior fronto-occipital fasciculus, optic radiation and vertical occipital fasciculus). Total lesion volume was compared between groups with a two-sided Brunner– Munzel test and related to detection accuracy by Spearman correlation. To test whether local lesion burden explained residual detection, the 33 parcels and tracts meeting a minimum-involvement criterion (mean lesion extent ≥ 10% across patients, or ≥ 10% in at least one patient) were retained. Lesion extent in each was correlated with detection accuracy (Spearman, FDR-corrected), and a mixed-effects model with region as a random effect and age and time since stroke as covariates tested the association between lesion extent and detection. Regional time series were averaged over all voxels of each parcel, including lesioned voxels. To test whether lesioned tissue contributed to the findings, the principal analyses were repeated after removing lesion-overlapping parcels, both as targets and as sources of each parcel’s regional mean MI. Three removal sets were used: the 32 cortical parcels lesioned by at least 10% in any patient, every parcel touched by a lesion in any patient, and each patient’s own lesioned parcels. In addition, the B+ vs. B− contrasts were repeated after regressing out total lesion volume or the mean extent of damage outside V1, alone or together with age, sex, time since stroke and lesion laterality (Supplementary Results).

### MRI acquisition

Data were acquired on a 3 T Siemens Prisma with a 64-channel head coil using the UK Biobank protocol [92, 93]. Resting-state fMRI used a multiband gradient-echo EPI sequence (2.4 mm isotropic; matrix 88 *×* 88 *×* 64; TR 735 ms; TE 39 ms; flip angle 52^*°*^; multiband factor 8; 490 volumes, ≈6 min). A T1-weighted MPRAGE was acquired at 1 mm isotropic (matrix 208 *×* 256 *×* 256; iPAT 2), together with field maps for distortion correction.

### Preprocessing and denoising

Functional data were processed with FSL: fieldmap-based distortion correction, motion correction (MCFLIRT), brain extraction (BET), grand-mean intensity normalization, band-pass filtering (0.008– 0.08 Hz) and boundary-based registration to the T1 image, which was normalized to MNI152 space with ANTs. Single-participant ICA components were classified with FIX using the UK Biobank training set and noise components were regressed out [94, 95]. Head motion was summarized as mean framewise displacement, which did not differ between groups and was unrelated to the unimodal–transmodal difference or to detection accuracy.

### Brain parcellations and time-series extraction

The brain was parcellated into 360 cortical regions [54] and 32 subcortical regions (scale II of Tian et al. [55]), giving 392 parcels. Denoised time series were averaged across voxels within each parcel and z-scored within participant. For group-level maps, the z-scored time series were concatenated across the participants of each group (HC, B+, B−) before MI estimation. All participant-level analyses used MI estimated separately for each participant over the 392 parcels, with each cortical parcel’s regional mean MI taken over the other cortical parcels. Where stated in Supplementary Results, group maps were computed as the mean of the participants’ individual MI matrices.

### Mutual information and regional mean MI

Mutual information (MI) is a core quantity in information theory that measures how much knowing one random variable reduces the uncertainty about another. In our context, it captures the extent to which the activity of one cortical region informs us about the activity of another, going beyond simple linear associations. Recent work by Luppi et al. [53], Varley et al. [96] and Santoro et al. [97] shows that such information-theoretic metrics offer complementary perspectives to conventional functional-connectivity approaches.

For two regional signals, *i* and *j*, mutual information is defined as

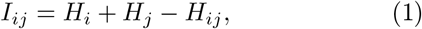

where *H*_*i*_ and *H*_*j*_ denote the Shannon entropies of the two signals, and *H*_*ij*_ is their joint entropy. It quantifies how much knowing one signal reduces uncertainty about the other. Under a bivariate Gaussian model, as assumed in standard functional-connectivity analysis,

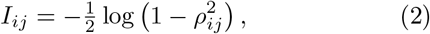

where *ρ*_*ij*_ is the Pearson correlation between the two regional time series. This estimator is a monotonic transform of |*ρ*| : it discards the sign of the correlation and weights strong correlations more heavily than the mean correlation does. Alternative estimators (rank-Gaussian copula, *k*-nearest-neighbor and binarized MI) and comparisons with signed and absolute mean correlation are reported in Supplementary Results. Each parcel’s regional mean MI was its average MI with all other analyzed parcels.

### Sensory-to-association axis and spin tests

Group-level cortical maps were compared with the S–A axis [43] by Spearman correlation across the 360 cortical parcels. Significance was assessed against a spatial null distribution of 5,000 rotations of the cortical sphere [98– 100]. Parcel values were reassigned by random rotation of the left hemisphere, with the right hemisphere mirroring it, which preserves the spatial autocorrelation of each map. *P*_spin_ is the one-sided proportion of rotations with a correlation at least as extreme as the observed one, in the direction of the observed correlation. Alternative references (cortical T1w/T2w ratio) and parcellations (Schaefer-200) are reported in Supplementary Results.

### Unimodal–transmodal differentiation

Parcels were assigned to the seven networks of Yeo et al. [60]. Unimodal cortex comprised the visual and somatomotor networks; transmodal cortex the frontoparietal and default-mode networks. For each participant, regional mean MI was averaged across unimodal and across transmodal parcels; their difference was the participant’s unimodal–transmodal differentiation. A somatomotor– transmodal difference was computed in the same way with the unimodal average restricted to the somatomotor network.

### Mixed-effects model

Regional mean MI values of unimodal and transmodal parcels were modeled with group, cortical pole (unimodal vs. transmodal) and their interaction as fixed effects and participant as a random intercept. The group contrasts reported in the Results are Wald tests on the group *×* pole interaction; that is, on the unimodal-minus-transmodal difference. Participant-level covariates entered as main effects cannot change this interaction, which is estimated within participants. In a separate model of the two patient groups, lesion volume, age, sex, time since stroke and lesion laterality were therefore allowed to modulate the difference (covariate *×* pole terms). Because these models treat each parcel’s value as a separate observation, a participant-specific random slope for cortical pole was also fitted, which tests the interaction against between-participant variability of the difference (Supplementary Results). Models were fitted by restricted maximum likelihood.

### Brain–behavior association and out-of-sample prediction

Across the 16 classified patients, the unimodal– transmodal difference was correlated with detection accuracy (Spearman’s *ρ*). The one-tailed *P* value was the proportion of 10,000 random re-pairings of the two variables yielding a correlation at least as large as the observed one. Partial associations adjusted for covariates were computed as Pearson correlations of rank residuals after regressing the covariates out of both variables, with one-tailed Freedman–Lane permutation *P* values (100,000 permutations). Out-of-sample prediction used leave-one-out cross-validation of a ridge regression of detection accuracy on the unimodal–transmodal difference (*λ* = 1, fixed, on the standardized predictor); cross-validated *R*^2^ was computed as 1*− SS*_res_*/SS*_tot_ over held-out predictions, and its *P* value as the proportion of 1,000 permutations of detection scores yielding a larger *R*^2^.

### Thalamic, seed and edge analyses

For each participant and each of the eight thalamic parcels of the Tian atlas (four per hemisphere), the parcel’s mean MI with unimodal parcels minus its mean MI with transmodal parcels was computed and compared between groups (one-tailed Brunner–Munzel tests, FDR-corrected). This difference was also correlated with detection accuracy as above. The cortical distribution of B+ *−* B− differences in pairwise MI with the surviving parcel was computed from MI matrices. In the whole-brain seed analysis, each of the 392 parcels was taken in turn as a seed. Its mean MI with somatomotor parcels minus its mean MI with transmodal parcels was computed per participant and compared between B+ and B− patients (one-tailed Brunner–Munzel, FDR-corrected). Pairwise MI between the thalamic parcel and each cortical seed that survived correction was then compared between groups (one-tailed against B−; two-sided for B+ vs. HC; FDR-corrected). Specificity was assessed by repeating the edge comparisons for the homologous contralesional thalamic parcel, and by ranking all 392 parcels by the separation of the patient groups in their coupling with the three cortical seeds under three criteria: i) the mean absolute Hedges’ *g* of the B+ vs. B− contrast across the edges; ii) the mean product of the HC vs. B− and B+ vs. B− effect sizes; and iii) the Stouffer-combined one-tailed B+ *>* B− significance across the edges.

### Statistical analysis

Group comparisons used the Brunner–Munzel test, which assumes neither normality nor equal variances. One-tailed tests were used where the direction was set a priori by the hypothesis stated in the Introduction: if the large-scale functional architecture also indexes non-conscious visual processing, B+ patients should show a more preserved cortical hierarchy than B− patients, approaching controls. The mixed-effects models test the same group *×* pole interaction with two-sided Wald tests, and in them the B+ vs. B− difference remained significant. *W* is signed so that a positive value indicates a larger value in the first-named group. Where the values of two groups did not overlap, leaving the Brunner– Munzel statistic undefined, the exact Mann–Whitney *P* is reported. One-sample comparisons against zero used the Wilcoxon signed-rank test. Effect sizes are Hedges’ *g*, with confidence intervals (CI) from the noncentral *t* distribution; correlations across participants are given with CI from the Fisher *z* transformation, with variance (1 + *ρ*^2^*/*2)*/*(*n −* 3 *− k*) for *n* participants and *k* covariates; and mixed-effects terms as the coefficient *b* (difference in the unimodal–transmodal gap, in MI units) with its Wald CI. All CI are at 95%, two-sided for two-tailed tests and one-sided, in the tested direction, for one-tailed tests. Multiple comparisons were corrected with the Benjamini–Hochberg false-discovery rate at *α* = 0.05 within each family of tests: the three group contrasts of each measure, the regions of each lesion analysis, the eight thalamic parcels, the 392 seeds and the tested thala-mocortical edges. Supplementary Results report sensitivity analyses for the MI estimator, parcellation and hierarchy reference, and for lesion volume, lesion-overlapping parcels and demographic and clinical covariates. Stability under participant sampling was assessed by boot-strap, leave-one-participant-out refits, group-label permutation and whole-pipeline relabeling, and within-run consistency by block bootstrap and intraclass correlation (Supplementary Results).

## CODE AVAILABILITY

Analyses were run in Python with NumPy, SciPy, pandas, statsmodels and pingouin (https://pingouin-stats.org). Functional data were preprocessed with FSL, including FIX and XTRACT (https://fsl.fmrib.ox.ac.uk), and ANTs (https://github.com/ANTsX/ANTs). Correlation matrices were estimated with nilearn (https://nilearn.github.io) and scikit-learn (https://scikit-learn.org). Spin tests used the ENIGMA Toolbox (https://github.com/MICA-MNI/ENIGMA). Figures were drawn with matplotlib and nibabel, and subcortical structures with subcortex-visualization (https://github.com/anniegbryant/subcortex_visualization).

## AUTHOR CONTRIBUTIONS

D.O., A.I.L. and M.T. conceived and developed the study idea; M.T. and H.B. designed the experiments; H.W. collected the data; M.D. preprocessed the data; D.O. analyzed the data; D.O., A.I.L., G.P. and M.T. wrote the original draft with valuable revisions by T.C., M.D., A.B. and M.P. All authors contributed to review and editing. M.T., G.P., and H.B. supervised the project. H.B. and M.T. are co-last authors. All authors approved the manuscript.

## References

[1] E. Pöppel, R. Held, and D. Frost, Residual visual function after brain wounds involving the central visual pathways in man, Nature 243, 295–296 (1973).

[2] L. Weiskrantz, E. K. Warrington, M. D. Sanders, and J. Marshall, Visual capacity in the hemianopic field following a restricted occipital ablation, Brain 97, 709–728 (1974).

[3] M. D. Sanders, E. K. Warrington, J. Marshall, and L. Weiskrantz, “Blindsight”: vision in a field defect, The Lancet 303, 707–708 (1974).

[4] L. Weiskrantz, Blindsight: A Case Study and Implications (Oxford University Press, Oxford, 1986).

[5] L. Weiskrantz, Consciousness Lost and Found: A Neuropsychological Exploration (Oxford University Press, Oxford, 1997).

[6] G. Rees, G. Kreiman, and C. Koch, Neural correlates of consciousness in humans, Nature Reviews Neuroscience 3, 261–270 (2002).

[7] F. Tong, Primary visual cortex and visual awareness, Nature Reviews Neuroscience 4, 219–229 (2003).

[8] D. Derrien, C. Garric, C. Sergent, and S. Chokron, The nature of blindsight: implications for current theories of consciousness, Neuroscience of Consciousness 2022, iab043 (2022).

[9] P. Stoerig and A. Cowey, Blindsight in man and monkey, Brain 120, 535–559 (1997).

[10] D. A. Leopold, Primary visual cortex: awareness and blindsight, Annual Review of Neuroscience 35, 91–109 (2012).

[11] J. Danckert and Y. Rossetti, Blindsight in action: what can the different sub-types of blindsight tell us about the control of visually guided actions?, Neuroscience & Biobehavioral Reviews 29, 1035–1046 (2005).

[12] M. A. Goodale and A. D. Milner, Separate visual pathways for perception and action, Trends in Neurosciences 15, 20–25 (1992).

[13] R. L. Whitwell, C. L. Striemer, D. A. Nicolle, and M. A. Goodale, Grasping the non-conscious: preserved grip scaling to unseen objects for immediate but not delayed grasping following a unilateral lesion to primary visual cortex, Vision Research 51, 908–924 (2011).

[14] J. E. LeDoux, M. Michel, and H. Lau, A little history goes a long way toward understanding why we study consciousness the way we do today, Proceedings of the National Academy of Sciences 117, 6976–6984 (2020).

[15] L. C. Sincich, K. F. Park, M. J. Wohlgemuth, and J. C. Horton, Bypassing V1: a direct geniculate input to area MT, Nature Neuroscience 7, 1123–1128 (2004).

[16] H. Bridge, O. Thomas, S. Jbabdi, and A. Cowey, Changes in connectivity after visual cortical brain damage underlie altered visual function, Brain 131, 1433–1444 (2008).

[17] M. C. Schmid, S. W. Mrowka, J. Turchi, R. C. Saunders, M. Wilke, A. J. Peters, F. Q. Ye, and D. A. Leopold, Blindsight depends on the lateral geniculate nucleus, Nature 466, 373–377 (2010).

[18] S. Ajina, F. Pestilli, A. Rokem, C. Kennard, and H. Bridge, Human blindsight is mediated by an intact geniculo-extrastriate pathway, eLife 4, e08935 (2015).

[19] A. Celeghin, M. Diano, B. de Gelder, L. Weiskrantz, C. A. Marzi, and M. Tamietto, Intact hemisphere and corpus callosum compensate for visuomotor functions after early visual cortex damage, Proceedings of the National Academy of Sciences 114, E10475–E10483 (2017).

[20] M. Kinoshita, R. Kato, K. Isa, K. Kobayashi, K. Kobayashi, H. Onoe, and T. Isa, Dissecting the circuit for blindsight to reveal the critical role of pulvinar and superior colliculus, Nature Communications 10, 135 (2019).

[21] H. E. Willis, M. R. Cavanaugh, S. Ajina, F. Pestilli, M. Tamietto, K. R. Huxlin, K. E. Watkins, and H. Bridge, Rehabilitating homonymous visual field deficits: white matter markers of recovery—stage 1 registered report, Brain Communications 6, fcae324 (2024).

[22] H. E. Willis, B. Caron, M. R. Cavanaugh, L. Starling, S. Ajina, F. Pestilli, M. Tamietto, K. R. Huxlin, K. E. Watkins, and H. Bridge, Rehabilitating homonymous visual field deficits: white matter markers of recovery— stage 2 registered report, Brain Communications 6, fcae323 (2024).

[23] I. Kletenik, M. A. Ferguson, J. R. Bateman, A. L. Cohen, C. Lin, A. Tetreault, V. S. Pelak, C. A. Anderson, S. Prasad, R. R. Darby, and M. D. Fox, Network localization of unconscious visual perception in blindsight, Annals of Neurology 91, 217–224 (2022).

[24] J. McFadyen, R. J. Dolan, and M. I. Garrido, The influence of subcortical shortcuts on disordered sensory and cognitive processing, Nature Reviews Neuroscience 21, 264–276 (2020).

[25] S. Ajina and H. Bridge, Blindsight relies on a functional connection between hMT+ and the lateral geniculate nucleus, not the pulvinar, PLoS Biology 16, e2005769 (2018).

[26] A. Sahraie, L. Weiskrantz, J. L. Barbur, A. Simmons, S. C. R. Williams, and M. J. Brammer, Pattern of neuronal activity associated with conscious and unconscious processing of visual signals, Proceedings of the National Academy of Sciences 94, 9406–9411 (1997).

[27] M. Tamietto, F. Cauda, L. L. Corazzini, S. Savazzi, C. A. Marzi, R. Goebel, L. Weiskrantz, and B. de Gelder, Collicular vision guides nonconscious behavior, Journal of Cognitive Neuroscience 22, 888–902 (2010).

[28] R. Goebel, L. Muckli, F. E. Zanella, W. Singer, and P. Stoerig, Sustained extrastriate cortical activation without visual awareness revealed by fMRI studies of hemianopic patients, Vision Research 41, 1459–1474 (2001).

[29] J. S. Morris, B. de Gelder, L. Weiskrantz, and R. J. Dolan, Differential extrageniculostriate and amygdala responses to presentation of emotional faces in a cortically blind field, Brain 124, 1241–1252 (2001).

[30] A. J. Pegna, A. Khateb, F. Lazeyras, and M. L. Seghier, Discriminating emotional faces without primary visual cortices involves the right amygdala, Nature Neuroscience 8, 24–25 (2005).

[31] M. D. Fox, A. Z. Snyder, J. L. Vincent, M. Corbetta, D. C. Van Essen, and M. E. Raichle, The human brain is intrinsically organized into dynamic, anticorrelated functional networks, Proceedings of the National Academy of Sciences 102, 9673–9678 (2005).

[32] B. J. He, A. Z. Snyder, J. L. Vincent, A. Epstein, G. L. Shulman, and M. Corbetta, Breakdown of functional connectivity in frontoparietal networks underlies behavioral deficits in spatial neglect, Neuron 53, 905–918 (2007).

[33] A. I. Luppi, H. M. Gellersen, L. Naci, A. Gozzi, and D. Bor, Unlocking spontaneous cognition in the age of data science, Nature Reviews Neuroscience 27, 591–593 (2026).

[34] D. Lyu, S. Naik, D. K. Menon, and E. A. Stamatakis, Intrinsic brain dynamics in the default mode network predict involuntary fluctuations of visual awareness, Nature Communications 13, 6923 (2022).

[35] I. Tavor, O. Parker Jones, R. B. Mars, S. M. Smith, T. E. Behrens, and S. Jbabdi, Task-free MRI predicts individual differences in brain activity during task performance, Science 352, 216–220 (2016).

[36] G. Pezzulo, M. Zorzi, and M. Corbetta, The secret life of predictive brains: what’s spontaneous activity for?, Trends in Cognitive Sciences 25, 730–743 (2021).

[37] A. R. Carter, S. V. Astafiev, C. E. Lang, L. T. Connor, J. Rengachary, M. J. Strube, D. L. W. Pope, G. L. Shulman, and M. Corbetta, Resting interhemispheric functional magnetic resonance imaging connectivity predicts performance after stroke, Annals of Neurology 67, 365–375 (2010).

[38] V. M. Saenger, A. Ponce-Alvarez, M. Adhikari, P. Hagmann, G. Deco, and M. Corbetta, Linking entropy at rest with the underlying structural connectivity in the healthy and lesioned brain, Cerebral Cortex 28, 2948–2958 (2018).

[39] C. Favaretto, M. Allegra, G. Deco, N. V. Metcalf, J. C. Griffis, G. L. Shulman, A. Brovelli, and M. Corbetta, Subcortical-cortical dynamical states of the human brain and their breakdown in stroke, Nature Communications 13, 5069 (2022).

[40] K. C. R. Fox, L. Shi, S. Baek, O. Raccah, B. L. Foster, S. Saha, D. S. Margulies, A. Kucyi, and J. Parvizi, Intrinsic network architecture predicts the effects elicited by intracranial electrical stimulation of the human brain, Nature Human Behaviour 4, 1039–1052 (2020).

[41] D. S. Margulies, S. S. Ghosh, A. Goulas, M. Falkiewicz, J. M. Huntenburg, G. Langs, G. Bezgin, S. B. Eickhoff, F. X. Castellanos, M. Petrides, E. Jefferies, and J. Smallwood, Situating the default-mode network along a principal gradient of macroscale cortical organization, Proceedings of the National Academy of Sciences 113, 12574–12579 (2016).

[42] J. M. Huntenburg, P.-L. Bazin, and D. S. Margulies, Large-scale gradients in human cortical organization, Trends in Cognitive Sciences 22, 21–31 (2018).

[43] V. J. Sydnor, B. Larsen, D. S. Bassett, A. Alexander-Bloch, D. A. Fair, C. Liston, A. P. Mackey, M. P. Milham, A. Pines, D. R. Roalf, J. Seidlitz, T. Xu, A. Raznahan, and T. D. Satterthwaite, Neurodevelopment of the association cortices: patterns, mechanisms, and implications for psychopathology, Neuron 109, 2820–2846 (2021).

[44] S. Park, K. V. Haak, S. Oldham, H. Cho, K. Byeon, B.-Y. Park, P. Thomson, H. Chen, W. Gao, T. Xu, S. Valk, M. P. Milham, B. Bernhardt, A. Di Martino, and S.-J. Hong, A shifting role of thalamocortical connectivity in the emergence of cortical functional organization, Nature Neuroscience 27, 1609–1619 (2024).

[45] J. Rittmo, N. Franzmeier, O. Strandberg, L. Chauveau, T. D. Satterthwaite, L. E. M. Wisse, N. Spotorno, H. H. Behjat, A. Dehsarvi, D. van Westen, T. E. Anijärv, Alzheimer’s Disease Neuroimaging Initiative, S. M. Landau, S. Palmqvist, S. Janelidze, E. Stomrud, R. Ossenkoppele, N. Mattsson-Carlgren, O. Hansson, and J. W. Vogel, Different functional connectivity gradients reflect aging and Alzheimer’s disease, Nature Neuroscience 29, 2835–2847 (2026).

[46] Y. Oh, Y. Ann, J.-J. Lee, T. Ito, S. Froudist-Walsh, C. Paquola, M. Milham, R. N. Spreng, D. Margulies, B. Bernhardt, C.-W. Woo, and S.-J. Hong, State-dependent signal flow hierarchy in the human cerebral cortex, Nature Neuroscience 29, 2848–2860 (2026).

[47] Z. Huang, G. A. Mashour, and A. G. Hudetz, Functional geometry of the cortex encodes dimensions of consciousness, Nature Communications 14, 72 (2023).

[48] A. I. Luppi, L. Uhrig, J. Tasserie, C. M. Signorelli, E. A. Stamatakis, A. Destexhe, B. Jarraya, and R. Cofré, Local orchestration of distributed functional patterns supporting loss and restoration of consciousness in the primate brain, Nature Communications 15, 2171 (2024).

[49] C. J. Whyte, M. J. Redinbaugh, J. M. Shine, and Y. B. Saalmann, Thalamic contributions to the state and contents of consciousness, Neuron 112, 1611–1625 (2024).

[50] M. J. Redinbaugh, J. M. Phillips, N. A. Kambi, S. Mohanta, S. Andryk, G. L. Dooley, M. Afrasiabi, A. Raz, and Y. B. Saalmann, Thalamus modulates consciousness via layer-specific control of cortex, Neuron 106, 66–75.e12 (2020).

[51] A. M. Bastos, J. A. Donoghue, S. L. Brincat, M. Mahnke, J. Yanar, J. Correa, A. S. Waite, M. Lundqvist, J. Roy, E. N. Brown, and E. K. Miller, Neural effects of propofol-induced unconsciousness and its reversal using thalamic stimulation, eLife 10, e60824 (2021).

[52] J. Tasserie, L. Uhrig, J. D. Sitt, D. Manasova, M. Dupont, S. Dehaene, and B. Jarraya, Deep brain stimulation of the thalamus restores signatures of consciousness in a nonhuman primate model, Science Advances 8, eabl5547 (2022).

[53] A. I. Luppi, F. E. Rosas, P. A. M. Mediano, D. K. Menon, and E. A. Stamatakis, Information decomposition and the informational architecture of the brain, Trends in Cognitive Sciences 28, 352–368 (2024).

[54] M. F. Glasser, T. S. Coalson, E. C. Robinson, C. D. Hacker, J. Harwell, E. Yacoub, K. Ugurbil, J. Andersson, C. F. Beckmann, M. Jenkinson, S. M. Smith, and D. C. Van Essen, A multi-modal parcellation of human cerebral cortex, Nature 536, 171–178 (2016).

[55] Y. Tian, D. S. Margulies, M. Breakspear, and A. Zalesky, Topographic organization of the human subcortex unveiled with functional connectivity gradients, Nature Neuroscience 23, 1421–1432 (2020).

[56] K. Mikellidou, J. W. Kurzawski, F. Frijia, D. Montanaro, V. Greco, D. C. Burr, and M. C. Morrone, Area prostriata in the human brain, Current Biology 27, 3056–3060.e3 (2017).

[57] M. Tamietto and D. A. Leopold, Visual cortex: the eccentric area prostriata in the human brain, Current Biology 28, R17–R19 (2018).

[58] H. E. Willis, I. B. Ip, A. Watt, J. Campbell, S. Jbabdi, W. T. Clarke, M. R. Cavanaugh, K. R. Huxlin, K. E. Watkins, M. Tamietto, and H. Bridge, GABA and glutamate in hMT+ link to individual differences in residual visual function after occipital stroke, Stroke 54, 2286–2295 (2023).

[59] H. E. Willis, J. Hameed, L. Starling, C. Kaltenbach, M. J. Moore, A. Khan, R. Maxwell, M. Tamietto, S. Ajina, and H. Bridge, Voxel-based lesion-symptom mapping localizes residual visual function in hemianopia, Journal of Neuroscience 45, e1263242024 (2025).

[60] B. T. T. Yeo, F. M. Krienen, J. Sepulcre, M. R. Sabuncu, D. Lashkari, M. Hollinshead, J. L. Roffman, J. W. Smoller, L. Zöllei, J. R. Polimeni, B. Fischl, H. Liu, and R. L. Buckner, The organization of the human cerebral cortex estimated by intrinsic functional connectivity, Journal of Neurophysiology 106, 1125–1165 (2011).

[61] A. Celeghin, A. Bagnis, M. Diano, C. A. Méndez, T. Costa, and M. Tamietto, Functional neuroanatomy of blindsight revealed by activation likelihood estimation meta-analysis, Neuropsychologia 128, 109–118 (2019).

[62] M. Tamietto, P. Pullens, B. de Gelder, L. Weiskrantz, and R. Goebel, Subcortical connections to human amygdala and changes following destruction of the visual cortex, Current Biology 22, 1449–1455 (2012).

[63] A. Bryant, Subcortex visualization: a toolbox for custom data visualization in the subcortex and cerebellum, Aperture Neuro 6, doi:10.52294/001c.165884 (2026).

[64] A. Schaefer, R. Kong, E. M. Gordon, T. O. Laumann, X.-N. Zuo, A. J. Holmes, S. B. Eickhoff, and B. T. T. Yeo, Local-global parcellation of the human cerebral cortex from intrinsic functional connectivity MRI, Cerebral Cortex 28, 3095–3114 (2018).

[65] J. B. Burt, M. Demirtaş, W. J. Eckner, N. M. Navejar, J. L. Ji, W. J. Martin, A. Bernacchia, A. Anticevic, and J. D. Murray, Hierarchy of transcriptomic specialization across human cortex captured by structural neuroimaging topography, Nature Neuroscience 21, 1251–1259 (2018).

[66] B. D. Fulcher, J. D. Murray, V. Zerbi, and X.-J. Wang, Multimodal gradients across mouse cortex, Proceedings of the National Academy of Sciences 116, 4689–4695 (2019).

[67] M. Massimini, F. Ferrarelli, R. Huber, S. K. Esser, H. Singh, and G. Tononi, Breakdown of cortical effective connectivity during sleep, Science 309, 2228–2232 (2005).

[68] V. A. F. Lamme and P. R. Roelfsema, The distinct modes of vision offered by feedforward and recurrent processing, Trends in Neurosciences 23, 571–579 (2000).

[69] M. Levinson, E. Podvalny, S. H. Baete, and B. J. He, Cortical and subcortical signatures of conscious object recognition, Nature Communications 12, 2930 (2021).

[70] A. K. Seth, Z. Dienes, A. Cleeremans, M. Overgaard, and L. Pessoa, Measuring consciousness: relating behavioural and neurophysiological approaches, Trends in Cognitive Sciences 12, 314–321 (2008).

[71] M. Wilke, K.-M. Mueller, and D. A. Leopold, Neural activity in the visual thalamus reflects perceptual suppression, Proceedings of the National Academy of Sciences USA 106, 9465–9470 (2009).

[72] J. Seo, D.-J. Kim, S.-H. Choi, H. Kim, and B.-K. Min, The thalamocortical inhibitory network controls human conscious perception, NeuroImage 264, 119748 (2022).

[73] Z. Fang, Y. Dang, A. Ping, C. Wang, Q. Zhao, H. Zhao, X. Li, and M. Zhang, Human high-order thalamic nuclei gate conscious perception through the thalamofrontal loop, Science 388, eadr3675 (2025).

[74] Y. Komura, A. Nikkuni, N. Hirashima, T. Uetake, and A. Miyamoto, Responses of pulvinar neurons reflect a subject’s confidence in visual categorization, Nature Neuroscience 16, 749–755 (2013).

[75] L. Pessoa, Understanding brain networks and brain organization, Physics of Life Reviews 11, 400–435 (2014).

[76] T. Livne, D. Kim, N. V. Metcalf, L. Zhang, L. Pini, G. L. Shulman, and M. Corbetta, Spontaneous activity patterns in human motor cortex replay evoked activity patterns for hand movements, Scientific Reports 12, 16867 (2022).

[77] A. Dimakou, G. Pezzulo, A. Zangrossi, and M. Corbetta, The predictive nature of spontaneous brain activity across scales and species, Neuron 113, 1310–1332 (2025).

[78] I.-C. Mundinano, D. M. Fox, W. C. Kwan, D. Vidaurre, L. Teo, J. Homman-Ludiye, M. A. Goodale, D. A. Leopold, and J. A. Bourne, Transient visual pathway critical for normal development of primate grasping behavior, Proceedings of the National Academy of Sciences 115, 1364–1369 (2018).

[79] H. Zhou, R. J. Schafer, and R. Desimone, Pulvinar-cortex interactions in vision and attention, Neuron 89, 209–220 (2016).

[80] R. W. Kentridge, C. A. Heywood, and L. Weiskrantz, Attention without awareness in blindsight, Proceedings of the Royal Society B: Biological Sciences 266, 1805–1811 (1999).

[81] A. Y. Nam, J. Shin, M. Tenney, B. Zhang, and Y. K. Hong, Somatosensory cortex shapes perceptual decision bias via the superior colliculus, Nature Communications (2026), doi:10.1038/s41467-026-78331-0.

[82] M.-H. Grosbras and T. Paus, Transcranial magnetic stimulation of the human frontal eye field facilitates visual awareness, European Journal of Neuroscience 18, 3121–3126 (2003).

[83] L. Chanes, A. B. Chica, R. Quentin, and A. Valero-Cabré, Manipulation of pre-target activity on the right frontal eye field enhances conscious visual perception in humans, PLoS ONE 7, e36232 (2012).

[84] J. W. Kurzawski, K. Mikellidou, M. C. Morrone, and F. Pestilli, The visual white matter connecting human area prostriata and the thalamus is retinotopically organized, Brain Structure and Function 225, 1839–1853 (2020).

[85] H. N. Halbertsma, J. A. Elshout, D. P. Bergsma, D. G. Norris, F. W. Cornelissen, A. V. van den Berg, and K. V. Haak, Functional connectivity of the precuneus reflects effectiveness of visual restitution training in chronic hemianopia, NeuroImage: Clinical 27, 102292 (2020).

[86] S. Ajina, K. Jünemann, A. Sahraie, and H. Bridge, Increased visual sensitivity and occipital activity in patients with hemianopia following vision rehabilitation, Journal of Neuroscience 41, 5994–6005 (2021).

[87] A. Barbot, A. Das, M. D. Melnick, M. R. Cavanaugh, E. P. Merriam, D. J. Heeger, and K. R. Huxlin, Spared perilesional V1 activity underlies training-induced recovery of luminance detection sensitivity in cortically-blind patients, Nature Communications 12, 6102 (2021).

[88] M.-H. Grosbras and T. Paus, Transcranial magnetic stimulation of the human frontal eye field: effects on visual perception and attention, Journal of Cognitive Neuroscience 14, 1109–1120 (2002).

[89] A. B. Chica, A. Valero-Cabré, P. M. Paz-Alonso, and P. Bartolomeo, Causal contributions of the left frontal eye field to conscious perception, Cerebral Cortex 24, 745–753 (2014).

[90] W. S. Kemball-Cook, E. R. Bader, A. D. Boro, E. N. Eskandar, and N. J. Killian, Electrical stimulation of the human pulvinar generates visual percepts, Brain Stimulation 19, 103231 (2026).

[91] S. M. Fleming, Towards an integrative neuroscience of metacognition, Nature Reviews Neuroscience 27, 789–806 (2026).

[92] K. L. Miller, F. Alfaro-Almagro, N. K. Bangerter, D. L. Thomas, E. Yacoub, J. Xu, A. J. Bartsch, S. Jbabdi, S. N. Sotiropoulos, J. L. R. Andersson, L. Griffanti, G. Douaud, T. W. Okell, P. Weale, I. Dragonu, S. Garratt, S. Hudson, R. Collins, M. Jenkinson, P. M. Matthews, and S. M. Smith, Multimodal population brain imaging in the UK Biobank prospective epidemiological study, Nature Neuroscience 19, 1523–1536 (2016).

[93] F. Alfaro-Almagro, M. Jenkinson, N. K. Bangerter, J. L. R. Andersson, L. Griffanti, G. Douaud, S. N. Sotiropoulos, S. Jbabdi, M. Hernandez-Fernandez, E. Vallee, D. Vidaurre, M. Webster, P. McCarthy, C. Rorden, A. Daducci, D. C. Alexander, H. Zhang, I. Dragonu, P. M. Matthews, K. L. Miller, and S. M. Smith, Image processing and quality control for the first 10,000 brain imaging datasets from UK Biobank, NeuroImage 166, 400–424 (2018).

[94] G. Salimi-Khorshidi, G. Douaud, C. F. Beckmann, M. F. Glasser, L. Griffanti, and S. M. Smith, Automatic denoising of functional MRI data: combining independent component analysis and hierarchical fusion of classifiers, NeuroImage 90, 449–468 (2014).

[95] L. Griffanti, G. Salimi-Khorshidi, C. F. Beckmann, E. J. Auerbach, G. Douaud, C. E. Sexton, E. Zsoldos, K. P. Ebmeier, N. Filippini, C. E. Mackay, S. Moeller, J. Xu, E. Yacoub, G. Baselli, K. Ugurbil, K. L. Miller, and S. M. Smith, ICA-based artefact removal and accelerated fMRI acquisition for improved resting state network imaging, NeuroImage 95, 232–247 (2014).

[96] T. F. Varley, M. Pope, M. G. Puxeddu, J. Faskowitz, and O. Sporns, Partial entropy decomposition reveals higher-order information structures in human brain activity, Proceedings of the National Academy of Sciences 120, e2300888120 (2023).

[97] A. Santoro, M. Neri, S. Poetto, D. Orsenigo, M. Diano, M. Gatica, and G. Petri, Charting higher-order models of brain function beyond pairwise interactions, Nature Communications 17, 9207 (2026).

[98] A. F. Alexander-Bloch, H. Shou, S. Liu, T. D. Satterthwaite, D. C. Glahn, R. T. Shinohara, S. N. Vandekar, and A. Raznahan, On testing for spatial correspondence between maps of human brain structure and function, NeuroImage 178, 540–551 (2018).

[99] R. D. Markello and B. Mišić, Comparing spatial null models for brain maps, NeuroImage 236, 118052 (2021).

[100] F. Váša and B. Mišić, Null models in network neuroscience, Nature Reviews Neuroscience 23, 493–504 (2022).

